# Prospects for recurrent neural network models to learn RNA biophysics from high-throughput data

**DOI:** 10.1101/227611

**Authors:** Michelle J Wu, Johan OL Andreasson, Wipapat Kladwang, William J Greenleaf, Eterna participants, Rhiju Das

## Abstract

RNA is a functionally versatile molecule that plays key roles in genetic regulation and in emerging technologies to control biological processes. Computational models of RNA secondary structure are well-developed but often fall short in making quantitative predictions of the behavior of multi-RNA complexes. Recently, large datasets characterizing hundreds of thousands of individual RNA complexes have emerged as rich sources of information about RNA energetics. Meanwhile, advances in machine learning have enabled the training of complex neural networks from large datasets. Here, we assess whether a recurrent neural network model, Ribonet, can learn from high-throughput binding data, using simulation and experimental studies to test model accuracy but also determine if they learned meaningful information about the biophysics of RNA folding. We began by evaluating the model on energetic values predicted by the Turner model to assess whether the neural network could learn a representation that recovered known biophysical principles. First, we trained Ribonet to predict the simulated free energy of an RNA in complex with multiple input RNAs. Our model accurately predicts free energies of new sequences but also shows evidence of having learned base pairing information, as assessed by *in silico* double mutant analysis. Next, we extended this model to predict the simulated affinity between an arbitrary RNA sequence and a reporter RNA. While these more indirect measurements precluded the learning of basic principles of RNA biophysics, the resulting model achieved sub-kcal/mol accuracy and enabled design of simple RNA input responsive riboswitches with high activation ratios predicted by the Turner model from which the training data were generated. Finally, we compiled and trained on an experimental dataset comprising over 600,000 experimental affinity measurements published on the Eterna open laboratory. Though our tests revealed that the model likely did not learn a physically realistic representation of RNA interactions, it nevertheless achieved good performance of 0.76 kcal/mol on test sets with the application of transfer learning and novel sequence-specific data augmentation strategies. These results suggest that recurrent neural network architectures, despite being naïve to the physics of RNA folding, have the potential to capture complex biophysical information. However, more diverse datasets, ideally involving more direct free energy measurements, may be necessary to train *de novo* predictive models that are consistent with the fundamentals of RNA biophysics.

**Author Summary:** The precise design of RNA interactions is essential to gaining greater control over RNA-based biotechnology tools, including designer riboswitches and CRISPR-Cas9 gene editing. However, the classic model for energetics governing these interactions fails to quantitatively predict the behavior of RNA molecules. We developed a recurrent neural network model, Ribonet, to quantitatively predict these values from sequence alone. Using simulated data, we show that this model is able to learn simple base pairing rules, despite having no *a priori* knowledge about RNA folding encoded in the network architecture. This model also enables design of new switching RNAs that are predicted to be effective by the “ground truth” simulated model. We applied transfer learning to retrain Ribonet using hundreds of thousands of RNA-RNA affinity measurements and demonstrate simple data augmentation techniques that improve model performance. At the same time, data diversity currently available set limits on Ribonet’s accuracy. Recurrent neural networks are a promising tool for modeling nucleic acid biophysics and may enable design of complex RNAs for novel applications.

## Background

RNA is involved in myriad functional roles in biological systems, from control of gene expression to splicing to protein synthesis, and serves as a key player in biotechnology tools, from RNA silencing to CRISPR/Cas9 to reengineered ribosomes.[1–8] These RNAs interact with each other and with other biomolecules in complex, dynamic ways that are highly dependent on the cellular environment in which they reside.[9,10] Understanding and controlling these processes requires a quantitative understanding of the energetics that govern these interactions. The classic nearest-neighbor model of nucleic acid energetics developed by Turner and colleagues is based upon careful parameterization of the energies of each possible motif formed.[11–13] However, this model makes strong assumptions about motif energies and interactions and relies on modest amounts of optical melting data.[11,14] Limited by the need for expert tuning and low measurement throughput, this model falls short in accurately predicting these interactions in many applications.[15,16]

Recently, technological developments have enabled high-throughput biophysical measurements of tens of thousands if individual RNAs, a rich source of data for refining models of RNA energetics. These techniques, which make use of repurposed Illumina sequencing tools, enable the measurement of RNA-RNA and RNA-protein affinities over millions of individual clusters on an RNA array in a wide variety of experimental conditions.[17–20] However, the complex relationship of these measurements to underlying energetic parameters makes it difficult to refine individual motif energies using these rich datasets. Thus, alternative models and optimization methods are needed to fully leverage these datasets for building quantitative models of RNA energetics.

Toward this end, advances in training neural network models have enabled extraction of relevant features from large, complex datasets. These deep learning models have recently propelled major improvements in many machine learning tasks, such as image classification, machine translation, and speech recognition.[21–24] They have also been successfully applied to modeling biological systems, although less work has been carried out in regression as opposed to classification tasks.[25–28] Among neural network architectures that have been successful in deep learning, recurrent neural networks (RNNs) have had a particularly large impact on processing temporal or sequence data, such as natural language or speech.[22,29,30] This architecture might be expected to naturally transfer to processing biological sequence data and has the potential to encode the level of complexity involved in making quantitative predictions of RNA interactions. Furthermore, such neural networks may be able to learn and predict non-nearest-neighbor effects and tertiary interactions that are not captured with classic physics-based nearest-neighbor models. Neural networks also offer the prospect of dramatic decreases in calculation speed for long RNA sequences or multi-RNA complexes, which currently incur computational expenses that scale with worse than polynomial time in sequence length.[31]

Here, we evaluate the effectiveness of a recurrent neural network, Ribonet, for predicting the energetics governing RNA interactions and recovering behavior consistent with biophysical principles. Specifically, our goal is to predict the energetics of RNAs in complex with other input RNAs at various concentrations. Using simulated free energy data, we demonstrate that, given enough training examples, Ribonet can capture base pairing information, despite having no *a priori* information about RNA biophysics and no explicit secondary structure information for the training molecules. In contrast, simulated affinity data, which involve free energy differences over multiple states and thus are a more complex function of secondary structure, were too indirect to allow Ribonet to learn this base-pairing level information. Nevertheless, these affinity-trained Ribonet models were accurate enough to facilitate design of novel RNA switches that modulate their affinity to a reporter in response to binding of an input. Further, we use transfer learning and data augmentation to train models on experimental RNA-RNA and RNA-protein affinity measurements, achieving good predictive performance on held-out test sets but failing to recover known principles of RNA biophysics. Together, these results suggest that RNNs are a promising architecture for use in predicting the energetics of RNA interactions and could potentially be extended to other complex biophysical measurements that are a function of sequence. However, more direct experimental measurements are necessary to enable training of physically realistic models.

## Results

### A recurrent network model of complex RNA interactions

RNNs are designed specifically for data that take the form of a sequence of elements and thus can be naturally applied to nucleic acid sequences. In fact, a direct parallel can be drawn between the mathematical operations applied in a classic partition function algorithm and those of a long short term memory (LSTM) unit, an architecture common in machine translation tasks.[23,32] In this mapping, the cell state stores the partition functions of subsequences, although the activation functions typically used in an LSTM are different from those in the Turner model parallel (Figure 1A & 1B).[33] To make our data compatible with these existing components of deep learning architectures, we encoded each measurement as a sequence of one-hot vectors to serve as input to an RNN. Each experimental measurement is characterized by the design sequence as well as the concentrations and sequences of zero or more input RNAs. The one-hot vectors representing the sequences of each RNA were multiplied by the log concentration of that species. Input RNA sequences were placed at the beginning of the input to the RNN, followed by the design RNA, with each strand separated by a zero vector (Figure 1C). The resulting sequence of vectors, representing each base in the sequence, was fed from 5′ to 3′ end as input to a 2-layer, 1024-unit LSTM RNN. The final states of each layer were then passed through two fully connected layers to aggregate the features extracted, resulting in a final numerical Δ*G* prediction (Figure 1D).

**Figure 1:**
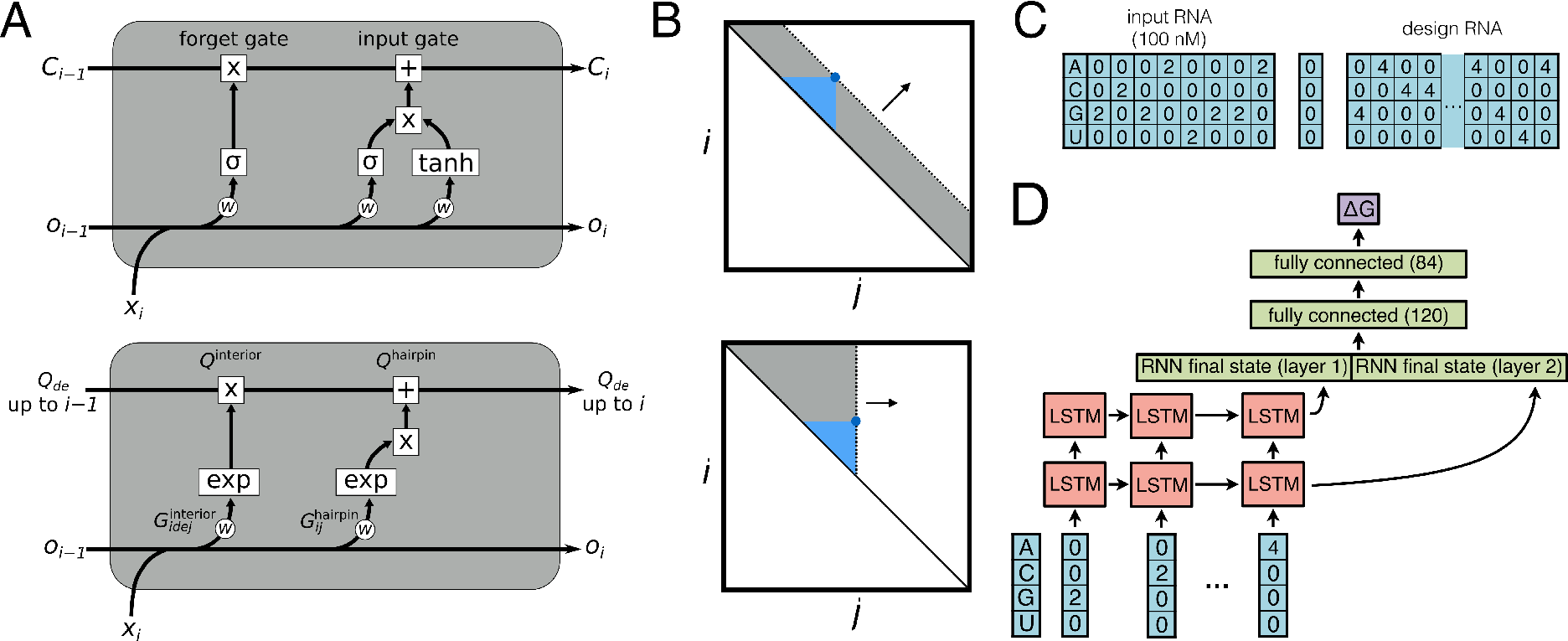
(A) Parallels can be drawn between the components of an LSTM unit (top) and those of a partition function calculation (bottom). The cell state (*C_i_*) can store the partition functions for subsequences (*Q_de_*), with the outputs (*o_i_*) containing the desired prediction value. The weights *w* in the LSTM contain the values that parameterize different motif energies (*G*). (B) The dynamic programming matrix can be computed horizontally rather than diagonally to account for the fact that the sequence is read in order, so that the 3′ end of the sequence is not needed at the start of the computation. (C) Each base in each sequence is encoded as an *n*-hot vector where *n* is the log of the RNA concentration. Input RNAs, followed by the design RNA, and, when relevant, the reporter RNA are taken in that order, and each strand is separated by a zero-vector. (D) The input is fed through a 2-layer LSTM RNN, followed by 2 fully connected layers.

### Exploring the expressivity of the RNN model on simulated free energies

As a first test of a recurrent neural network representation of RNA folding, we evaluated the expressivity of Ribonet on simulated energetic measurements from the existing nearest-neighbor model. Success in this task would demonstrate that an RNN can mimic a model that is constrained by known biophysical principles, suggesting the architecture may be suitable for modeling RNA energetics.

We began by training on free energy values for each complex relative to the fully unfolded and dissociated RNA, the most direct energetic characterization of an RNA fold (Figure 2A). Such data can be obtained empirically through optical melting experiments, albeit not at the throughput possible for other, less direct energetic measurements. We trained the model on simulated folding free energies for complexes consisting of randomly generated designs in 15 different conditions chosen to match available experimental measurements (1, 10–20 in Table 2). Designs were generated by introducing the reverse complements of the input RNAs into a random sequence at random positions (see Methods for details). For each design, the free energy was computed over a full ensemble of simulated secondary structures, as would best mimic an actual experimental scenario. As a standard of comparison for this dataset, two different parameter sets of the Turner model available in the NUPACK software gave calculations with root-mean-square error (RMSE) of 7.65 kcal/mol, giving a rough estimate of the error resulting from these parameter estimates. A 1-nearest neighbor model, which predicts based on the closest measurement in the training set, yielded an RMSE of 18.1 kcal/mol, setting a rough upper on our performance (Figure S1). To determine how much data are necessary to parameterize this model, we trained on datasets of varying size from 10,000 to 500,000 sequences. We found that 500,000 training sequences were needed in training to avoid overfitting (Figure 2B). Using this training set, we were able to achieve excellent predictive performance of 5.46 kcal/mol, better than the error between two Turner parameter sets (Figure 2C).

**Figure 2:**
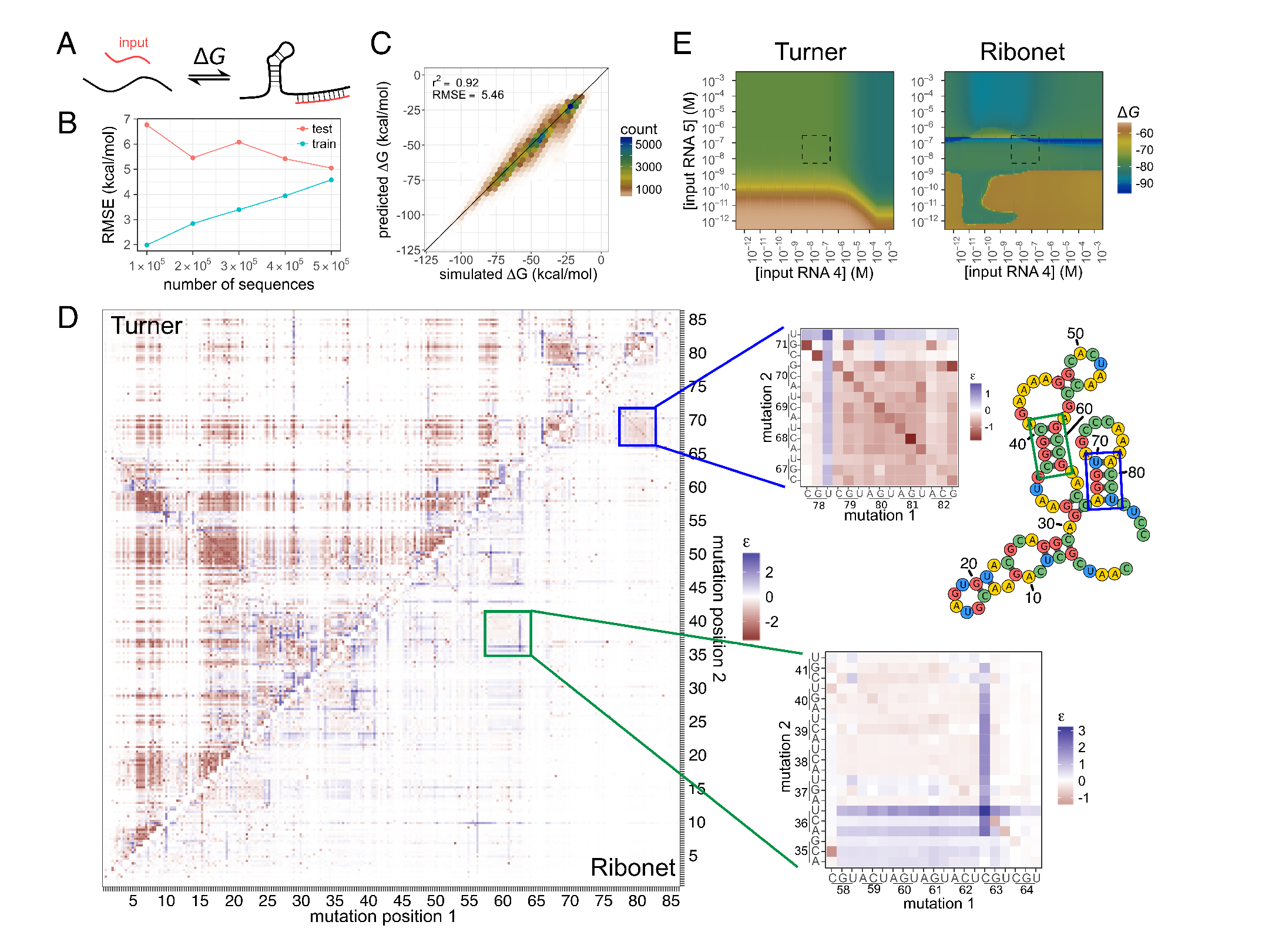
(A) The free energy computed is the difference between the folded complex and unfolded, dissociated individual strands. (B) For the free energy model, the learning curve for varying numbers of simulated sequences reveals that training and test performance converges at around 500,000 sequences. (C) For the largest dataset, the model achieves predictions of affinity with an RMSE of 5.46 kcal/mol. (D) For the free energy model, predictions were made for free energy of folding for a set of double mutants (lower right) and compared to those from the Turner model (upper left). To highlight the effects of double mutants, these are shown as epistatic effects (e), which is the predicted free energy of the double mutant minus those for the two corresponding single mutants. Diagonal signals (inset) suggest that the model detects compensatory rescue of base pairs. The secondary structure suggested by the helices is shown on the middle right. (E) Extrapolating to predictions outside the training conditions results in an irregular, unrealistic pattern (right), notably different from the simulated behavior (left). The boxed region highlights the range of concentrations seen in the training data, showing similar simulated and predicted behavior.

As a test of whether the trained model was able to represent detailed secondary structure information, we evaluated the model on all double mutants of a randomly generated sequence distinct from the training data (Figure 2D). This analysis revealed that the RNN is appropriately sensitive even to small changes in the input sequence. Moreover, it detects base pairing effects, seen as diagonal segments on a heat map of double mutants; two mutations can restore the formation of a Watson-Crick base pair broken by single mutations (Figure 2D, insets). However, not all such base-pairing signals computed in the Turner model simulations (Figure 2D, top left) were captured by Ribonet and signals predicted by the RNN model were typically weaker, indicating less sensitivity to base-pairing than an explicit representation. Finally, we made predictions for input concentrations outside the range seen in the training data, to test if the model learned biophysically reasonable concentration effects. This free energy model produced an irregular pattern that does not match that of the simulated pattern over concentration space (Figure 2E). This suggests that, given enough data, the RNN model is expressive enough to capture the behavior of the Turner model of RNA energetics but cannot correctly extrapolate outside the concentration range seen in training.

### Applying Ribonet to simulated affinity measurements

To more closely replicate how existing high-throughput experimental datasets might enable Ribonet training, we then generated a simulated dataset of predicted affinities to a reporter RNA molecule. For each sequence, we simulated the RNA-reporter affinities for 15 different conditions (1, 10–20 in Table 2). Sequences were randomly generated as in the folding free energy simulations, but affinities were computed by finding the difference in free energy between the complexes with and without the reporter molecule (Figure 3A, see Methods for details). As a comparison point, two different Turner parameter sets differ by an RMSE of 1.08 kcal/mol over the same sequences, while a simplistic 1-nearest neighbor model based on sequence similarity achieves an RMSE of about 3.3 kcal/mol (Figure S2). A learning curve revealed that more than 300,000 sequences were necessary to avoid overfitting the model (Figure 3B). Performance of 1.54 kcal/mol was achieved even with only 10,000 sequences (Figure S3A), less than the number of data collected in high-throughput experiments. The model trained on 500,000 sequences achieved even better predictive performance, with a test RMSE of 1.15 kcal/mol (Figure 3C). Although this model required fewer data to reach convergence than the absolute free energy study above, the resulting model does not exhibit base-pairing effects, as was the case in the free energy-trained model, and again did not extrapolate well to concentrations outside those seen in training (Figure 3D & E). We also tested a model trained on randomly generated input RNA sequences, rather than a fixed set of input sequences, to determine if a more diverse set of data would provide enough additional biophysical information. While these models did not perform better in terms of display of base pairing signals in double mutant tests or extrapolation to out-of-range input RNA concentrations, they did make more accurate predictions for complexes involving input RNAs not seen in training (Figure S3B & S3C). These tests suggest that Ribonet is able to capture affinity information with strong predictive power but without accurately capturing base-pairing effects or the functional form of affinity expected from statistical mechanics.

**Figure 3:**
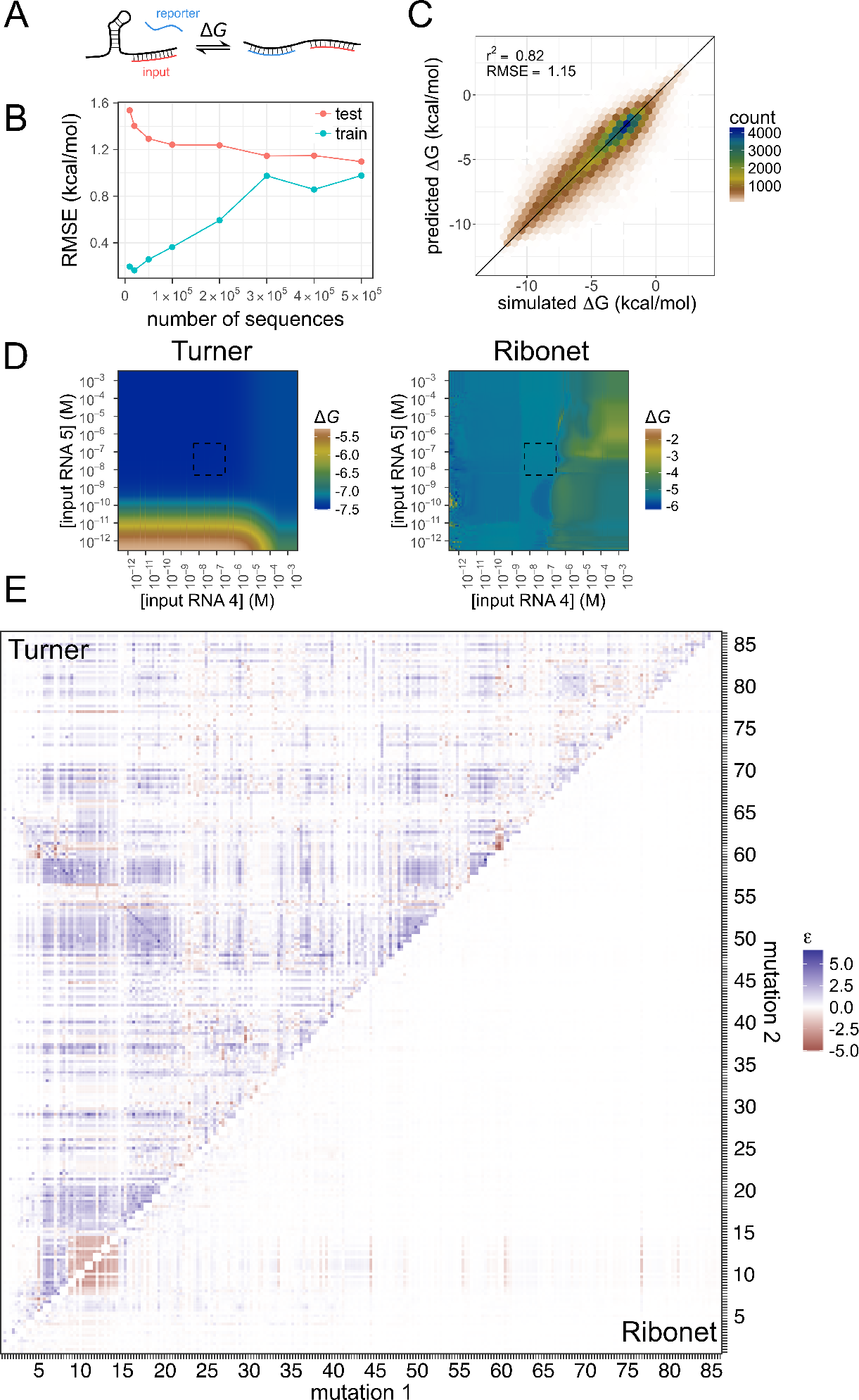
(A) The affinity is computed as the difference in free energy between the complexes with and without the reporter (blue). (B) For the simulated affinity data, a learning curve demonstrated that 300,000 sequences are necessary for convergence of training and test performance, fewer than for the free energy data. (C) For the largest dataset, the model achieves an RMSE of 1.15 kcal/mol. (D) The model (right) shows more realistic, smooth concentration dependence than the free energy model, even though the observed pattern differs greatly from the simulated one (left). The dotted box highlights the concentration range seen in the training set. (E) While the simulated affinities for a set of double mutants shows diagonal helix signals (upper left), this is not seen in the RNN predictions (lower right). The values shown are the predicted energy for the double mutant minus those for the two corresponding single mutants.

Despite its inability to recover physically realistic aspects of RNA folding, the accuracy of Ribonet for estimating affinities may still be useful for guiding design. We used a Monte Carlo algorithm to design an RNA sensor for an input RNA molecule, using the RNN as a scoring function and the Turner calculations as gold standard (Figure 4A, see Methods for details). In sequence optimization, we allowed both mutations and shifts of the input and reporter binding sites (Figure 4B). While shifts were accepted more often, mutations often preceded score increases, suggesting they are essential to sampling sequence space effectively (Figure 4C). We found that almost all designs improved over the initial sequence, as predicted by the Turner model, with the best designs achieving a fold change in affinity of almost 10^5^ (Figure 4D). The RNN-predicted fold changes agreed well with those simulated by the Turner model. These simulation results suggest that a model trained on sufficiently large datasets can represent the behavior of the RNA well enough to design new sequences from scratch. However, design became less effective with a model trained on a smaller dataset (Figure 4E). Further, it failed for design of sensors for input RNAs not seen in the training dataset (Figure S4A), even using the model trained on random input sequences (Figure S4B).

**Figure 4:**
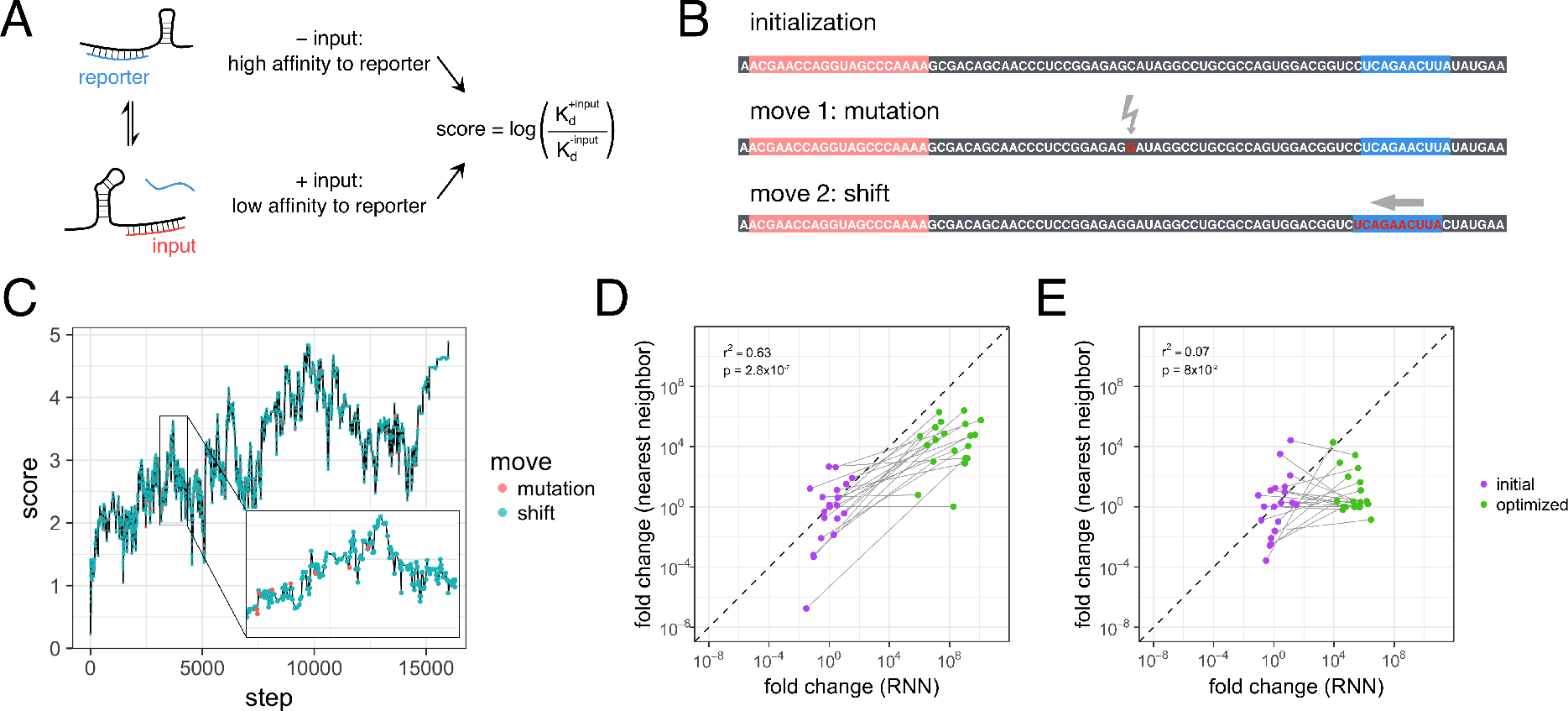
(A) For design, sequences were optimized using a score based on the ratio of affinities between the conditions with and without the input molecule. (B) Our design method initializes sequences with the reverse complements of the input and reporter sequences. Allowed moves include mutations and shifts of the reverse complements. (C) Our design method enables optimization of the RNN-predicted score. While most accepted moves are shifts (blue), significant jumps in score are often preceded by mutations (red). (D) The ground truth fold change between the two states, that predicted by the Turner model, increases significantly from initialization (purple) to after optimization (green), reaching a fold change as high as 10^6^. Lines connect the start and end sequences from the same design run. Predicted and ground truth values correlate well (*r*^2^ = 0.63), and a paired one-sided *t-*test shows that the optimized fold changes are significantly greater than the initial ones (*p*=2.8×10^−7^). (E) For the model trained on 10,000 sequences, the improvement is not as consistent, not reaching statistical significance (*p*=0.08), but most runs still yield an improvement in nearest-neighbor predicted fold change, our ground truth in these simulations.

### Evaluating Ribonet on experimental affinity measurements

Having gained an understanding of the nuances of Ribonet’s model architecture in relation to existing computational models, we trained the model on a rich dataset of affinity measurements. The Eterna massive open laboratory enables citizen scientists to design complex, multi-state RNAs using an intuitive graphical user interface.[20] Details of player strategies and performance are being reported in a separate manuscript; a brief summary follows. Various design challenges were posed as in-game puzzles with secondary structure constraints over multiple conditions defined by the concentrations of one or more input RNA strands. Players started with switch puzzles, where the aim was to design a molecule whose binding to a reporter RNA is dependent upon the presence or absence of a single input RNA. Next, players designed logic gates, which modulate their reporter binding based on the presence or absence of two input RNAs. Lastly, players moved to designing RNAs with an analog response, tuning the reporter affinity based on the ratio of concentrations of two to three input RNAs. Over 65,000 designs were made for these four puzzle types, collected over 9 different rounds (Table 1). While the library of designs contains a fair number of highly similar sequence clusters, it still represents a diverse collection of sequences targeting these 3 different types of puzzles (Figure 5A). Each sequence was synthesized on an RNA array, and its affinity to a fluorescent reporter RNA was measured by quantifying the fluorescence over varying reporter concentrations using a repurposed Illumina sequencing machine (see Methods for more details).[17] Each design was measured in up to 19 different input conditions, resulting in over 600,000 distinct affinity measurements.

**Figure 5:**
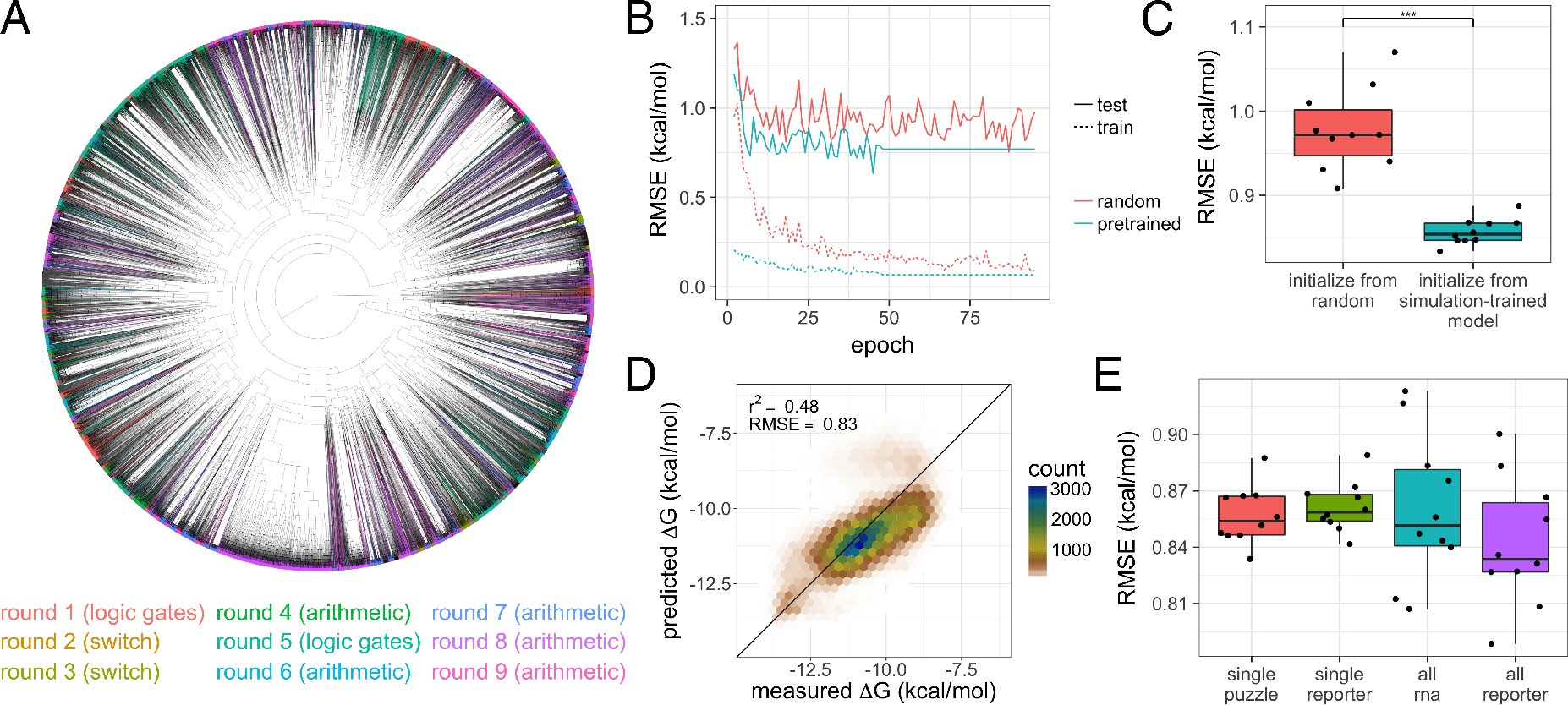
(A) Hierarchical clustering of all sequences designed over 9 rounds of collection show many tight clusters of similar sequences. (B) Initializing weights from a model pretrained on simulated data allows the loss to converge more quickly and to a lower final train and test RMSE than the random initialized model. The pretrained model shown was only trained for an additional 50 epochs, and the final loss is extended horizontal for ease of comparison. (C) Pretraining the model significantly improves performance (one-sided *t*-test, *p* = 6.46×10^−6^). (D) With transfer learning, the model is able to achieve performance of 0.83 kcal/mol. (E) Test results are shown for 10 replicate training experiments for each model type. No statistically significant differences are seen across models. All RMSEs are for only test set points with Levenshtein distance of at least 5 from all sequences in the training set.

**Table 1:**
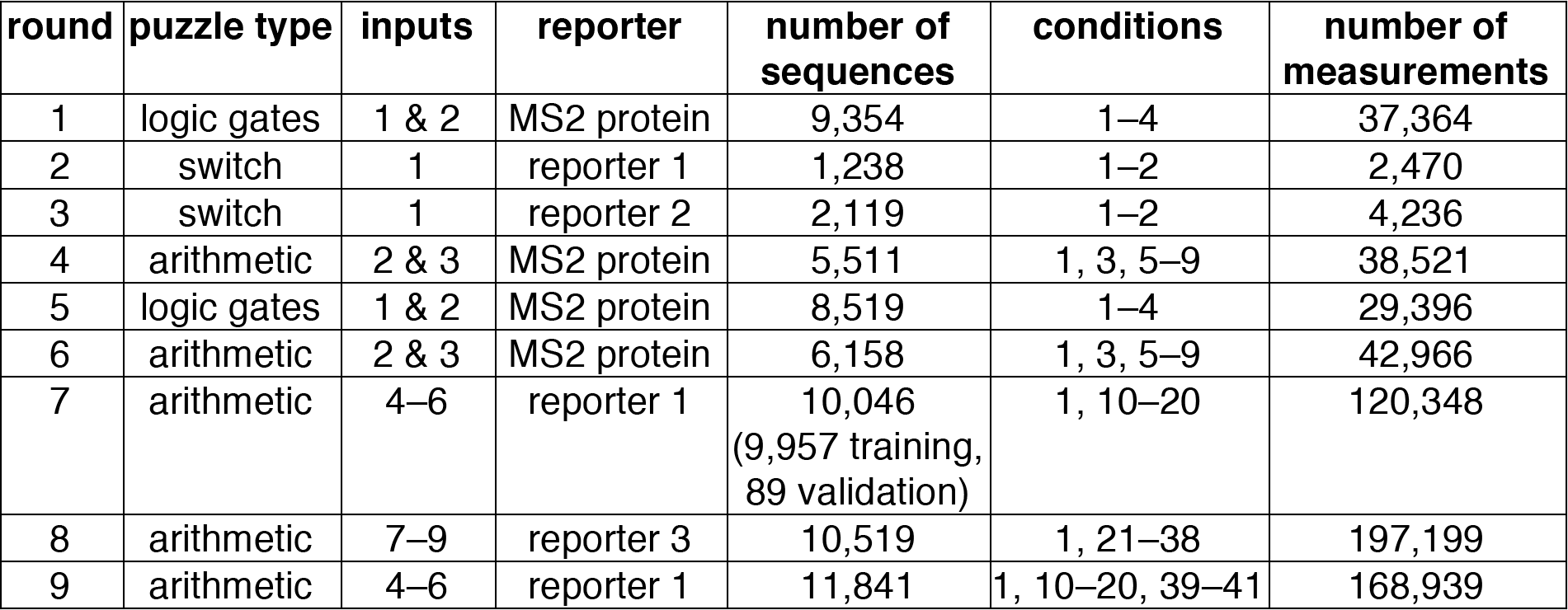
Summary of data collection rounds. Specific sequences and concentrations for each condition are available in Table 2 & 3. The number of measurements is less than the product of the number of sequences and the number of conditions due to filtering of low quality measurements.

Notably, these data are limited in range due to experimental constraints and contain less diverse sequences than the random simulated data (Figure 5A). To set a baseline for how well Ribonet might learn from realistic experimental datasets, we applied constraints to our simulated data to more closely match the experimental data. First, we filtered out points in the training data outside the experimentally measurable range, which reduced the 500,000 sequence dataset by about 5-fold. Test performance for data within the filtered range was good, with an RMSE of 0.92 kcal/mol, although worse than a similarly sized, unfiltered dataset, with an RMSE of 1.24 kcal/mol over values with nearly five-fold higher variance (Figure S5A). For test points outside the window, Δ*G* values were predicted to be near the edge of the measurable range (Figure S5B). This pileup is the expected behavior, because the model is unlikely to predict values beyond the range seen in the training data; however, the result underscores the inability of the model to learn physically reasonable extrapolations. In addition, we tested the model on simulated Δ*G*s for the sequences probed in available experiments instead of randomly generated sequences. The performance was far poorer than that for a comparably-sized randomly generated dataset, most likely due to the lower diversity of sequences (Figure S5C).

We then moved to experimental data, starting with a Ribonet model trained only on data from a single type of puzzle, with a fixed reporter sequence and a fixed set of three input RNAs (round 7 in Table 1). We used separate rounds of design and experiments for our training and test sets, with the test set having 3 additional experimental conditions not seen in the test set. As a lower bound on expected performance, the RMSE between technical replicates, which is a lower bound on our performance, is 0.39 kcal/mol (Figure S6A). As an upper bound, the RMSE for a 1-nearest neighbor model based on sequence similarity is 1.20 kcal/mol (Figure S6B, see methods for details). This bound is significantly tighter than that for simulation studies above due to the low sequence diversity of the designs and experimental measurement limits, setting a stringent bound for the performance of Ribonet. Further, the classic Turner model of RNA folding predicts these data poorly, with an RMSE of 2.73 kcal/mol (Figure S7). Training on 120,348 measurements over 10,046 sequences, we found that the model typically converged after about 50 epochs and significantly overfit to the training data, as evidenced by a large gap between training and test accuracy (Figure 5B). In our analysis of test set prediction results, we applied a Levenshtein distance threshold to the closest sequence in the training set to ensure that we did not include measurements unfairly similar to those in the training data (see Methods for details). For this model, the best model reaches an RMSE of 0.91 kcal/mol (Figure S8A). including predictions for experimental conditions not seen in the training data (Figure S9).

Because we had already trained Ribonet on simulated data from the Turner model, we hypothesized that performing transfer learning from the simulated model would accelerate and improve training. Thus, we trained models initialized with weights from the 500,000-sequence trained affinity model and compared those to randomly initialized weights. The training loss converged much more quickly, as expected, and also to a better test accuracy of 0.83 kcal/mol (Figure 5B). While the models were still overfit, as evidenced by a large gap between test and training accuracy, the performance was not only significantly better but also much more consistent across runs (Figure 5C). The preinitialized runs reached a median RMSE of 0.85 kcal/mol, with the best model reaching 0.83 kcal/mol (Figure 5D). For the rest of the models described, the transfer learning approach was used to initialize weights.

We then expanded our RNN models to increasing levels of generality, beyond the “single puzzle” setup used as an initial test case. As models become more general, the number of distinct puzzles and the associated amount of data available to train the model increases. However, the space of interactions that the model must represent also increases as the data involve more diverse inputs and reporters. For example, 29,542 additional sequence designs involve binding of an MS2 viral coat protein to specific RNA binding sites, but this protein-RNA interaction is captured through a new energetic parameter instead of those of RNA-RNA binding in designs tested previously. We tested three additional levels of model generality. “Single reporter” models generalized over multiple puzzles with the same reporter sequence, “all RNA” models included all measurements with RNA reporters over many puzzles and reporters, and “all reporter” models further included measurements with protein-based reporters. Empirically, we found that the addition of more complex RNA reporter data had no effect on performance (Figure 5E & S8), but predictive performance for protein reporters was poor, with RMSE of 1.06 kcal/mol (Figure S10). Our results suggest that training a more general model can allow for broader usability of the resulting model without compromising the predictive performance. In the case of including non-RNA reporters, however, additional information may need to be encoded to help the RNN understand how the reporter interacts with the design molecule.

### Data augmentation to enhance model performance

Our analysis on simulated data suggested that a dataset of tens of thousands sequences, such as that used in each of our models is not sufficient to obtain optimal performance. To better understand the data dependence of this RNN for real data, we trained the model on varying sized subsets of the data to build a learning curve.[34] To generate these datasets, we clustered the sequences based on Levenshtein distances and cut the tree to form 10 subsets of unequal size but similar distance between clusters (see Methods for more details). Training on subsets of data containing increasing numbers of clusters revealed that a strong downward trend in RMSE continues up to the maximum dataset size (Figure 6B). These learning curves suggest that the performance of Ribonet remains strongly data-limited.

**Figure 6:**
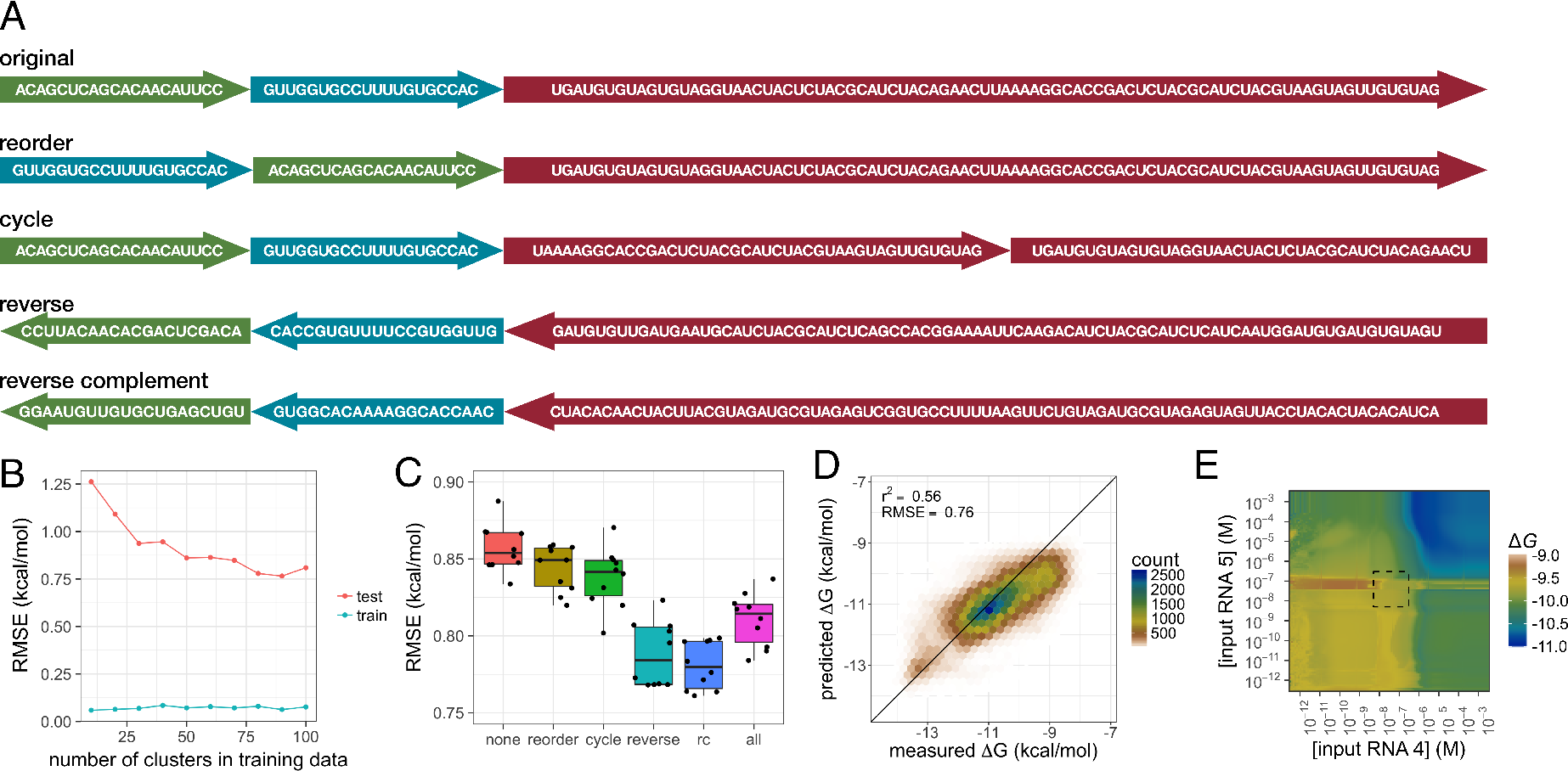
(A) Four different transformations were evaluated for data augmentation. The visualization shows the design sequence in red and two input sequences in green and blue. (B) A learning curve for the “single puzzle” reveals that the model is overfit, with test set RMSE exceeding train set RMSE, and lies in a high variance regime. (C) Augmentation of the training set significantly improves performance for the single puzzle model, with one-sided *t*-test *p-*values of 3.4×10^−2^ for reorder, 1.2×10^−2^ for cycle, 1.3×10^−7^ for reverse, 9.4×10^−10^ for reverse complement (RC) and 2.8×10^−6^ for all augmentation methods. (D) With reverse complement augmentation, the model achieves an RMSE of 0.76 kcal/mol. (E) In concentration extrapolation tests, the best model exhibits relatively smooth behavior compared to previous simulated models. The dotted box highlights the concentration range seen in training.

Data augmentation is a method commonly used in deep learning to increase the effective dataset size.[21,35,36] Although the techniques for augmentation are most well-established for image data, we hypothesized that the same ideas might be expanded to our datasets. We tested four approaches to augmenting our dataset (Figure 6A). First, we introduced additional copies of each data point for each possible ordering of the input sequences, since the chosen ordering is arbitrary. Secondly, we added a copy of each data point with a circular permutation of the design sequence. We have observed that top solutions of Eterna players are often circular permutations of each other (unpublished work). Thus, we hypothesize that this transformation may produce designs with similar behavior. Thirdly, reflection of input sequences has been used in machine translation to transform input data, despite the lack of semantic meaning in such data.[23] We introduced an additional copy of each data point with each sequence reversed, passed from 3′ to 5′ end instead of 5′ to 3′. This augmentation has the potential to increase signal for base pairing, as the equivalent helices can still form in a reflected secondary structure, even though the energetics of these alternative helices are different according to the classical Turner model.[11] Lastly, in a similar effort to encode additional base pairing information, we reverse complement each sequence to create an additional copy of each data point. To evaluate the effect of these augmentations on model performance, we used the “single puzzle” model, again using 10 trials with different random seeds to assess the variance. Although only the reordering approach worked in simulation (Figure S11), we observed a significant improvement in the performance for experimental data with the models trained on every type of augmented data, with *p*-values of 3.4×10^−2^ for reordering of inputs, 1.2×10^−2^ for circular permutations, 1.3×10^−7^ for reversed sequences, 9.4×10^−10^ for reverse complements, and 2.8×10^−6^ for all methods applied simultaneously (Figure 6C). The reverse complement augmentation method was able to achieve an RMSE of as low as 0.76 kcal/mol, compared to 0.83 kcal/mol without augmentation (Figure 6D). Performance with varying distance cutoffs is shown in Figure S12. These results suggest that simple transformations are able to encode additional information in model training, and reverse complementation is particularly effective, potentially by introducing additional base pairing information. These types of strategies may enable models to learn more biophysically relevant information from less data.

The best model trained with data augmentation still does not learn base pairing information in a test on double mutants (Figure S13). However, the behavior over wide concentration ranges is relatively smooth, with the exception of a discontinuity at 100 nM of the second input RNA (Figure 6E). This discontinuity may be a result of this being a frequent concentration choice in our set of experimental conditions. Overall, these tests suggest that Ribonet has the potential to produce biophysically realistic representations over concentration extrapolation, despite containing no explicit constraints to force this behavior. It remains unclear whether Ribonet can learn base-pairing information from simulated or experimental affinity data.

## Discussion

Here, we describe a novel RNN-based method, Ribonet, for predicting the affinity of RNA-RNA interactions in the context of different environmental conditions. We show that it exhibits the expected base pair effects when trained with simulated folding free energy data generated using a classic Turner model, despite the lack of *a priori* information about RNA folding. When instead trained with simulated affinity data, models lack this explicit biophysical information but still perform well enough to facilitate design of an RNA sensor. We successfully apply transfer learning to train Ribonet on data generated through the Eterna platform and high-throughput array experiments. We find that the models generally benefit from more diverse data but are unable to learn from a naïve representation of protein reporters. Finally, we introduce data augmentation techniques specific to nucleic acid sequence data that significantly improve performance.

This work represents one of the first applications of deep learning to a regression task in biology.[37,38] While we were able to achieve fairly good performance, as compared to prior biophysical models or 1-nearest neighbor models, we have shown that we are currently limited by the size of our dataset, a common theme in machine and deep learning.[39,40] Despite the use of high-throughput design and measurement, one limitation of our methodology is that, when design constraints become difficult to satisfy, Eterna players tend to produce additional submissions by modifying existing designs. This often results in clusters of sequences that are highly similar, differing only by single mutations or base pair swaps. Although these near redundancies have the potential to improve sensitivity to small changes and provide useful base pairing information, they also result in a smaller effective dataset size. Our data augmentation approach partially mitigated this effect, but further efforts would benefit from more diverse designs as well as development of experimental methods able to probe a much larger variety of input RNAs and reporter RNAs. Additional work is necessary to develop computational methods to enable us to take full advantage of the specific features of this dataset and to understand why the reverse complement augmentation aids accuracy in experimental but not simulated datasets.

Through our simulation studies, we also found that the indirect nature of the affinity measurements prevented learning of explicit information about base pairing, a feature that is fundamental to RNA interactions. Nevertheless, free energy values were sufficient for learning low level biophysical information, suggesting high-throughput approaches for making these kinds of measurements would open the door for more biophysically realistic models of RNA energetics.[41,42]

In addition, the architecture chosen here does not take advantage of existing knowledge about RNA folding. Many successful applications of deep learning have taken this approach of learning from scratch.[21,23] Although we have demonstrated here that the RNN is able to extract base pairing information *de novo*, it fails to do so from simulated or experimental affinity data, which is a more complex function of secondary structure interactions compared to free energies. While Ribonet-facilitated design was possible in simulation, it required an order of magnitude more data than currently available even with high-throughput experiments. Adding constraints to the weights or explicitly encoding additional information may accelerate model training or improve test performance.[43,44] Further work is necessary to optimize deep learning model architectures for specific biological systems.

Machine learning methods such as those described here may form the foundation for computational tools for RNA design. Given the ability to quantitatively predict RNA-reporter affinities, optimization algorithms could be applied to design RNAs that sense and interact with other biomolecules in their environments, as demonstrated by our design tests in simulation.[45–47] This level of predictive performance would enable us to precisely design interactions in biological systems, including those central to modern biotechnological tools.

## Materials and Methods

### Eterna massive open laboratory

The Eterna online platform (https://www.eternagame.org) enables citizen scientists to design RNA molecules with complex, multi-state folding behavior. The game allows for the specification of design challenges, or puzzles, with secondary structure constraints in multiple conditions defined by the concentrations of various input RNAs. Players can view folds and energies predicted by ViennaRNA 1.8.4 [48], Vienna 2.1.9 [49], and NUPACK 3.0.4. [31,47]

### RNA array experiments

Clonal clusters of RNA were generated on an Illumina sequencing flow cell as previously described.[17,18] Binding curves were collected for each cluster by incubating with input RNA oligos at defined concentrations and progressively higher concentrations (by factors of 2) of fluorescently labeled reporter oligo or recombinant SNAP-tag-labeled MS2 coat protein (Tables 1–3). The starting concentration was between 0.09 and 0.75 nM, depending on the experiment and the reporter, and the final concentration was 3000 nM. The fluorescent cluster images were aligned to sequencing data and quantified as previously described.[17,18] The fluorescent signal for each cluster, *F,* was fit to a simple binding curve:

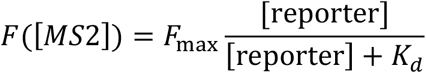

**Table 2:**
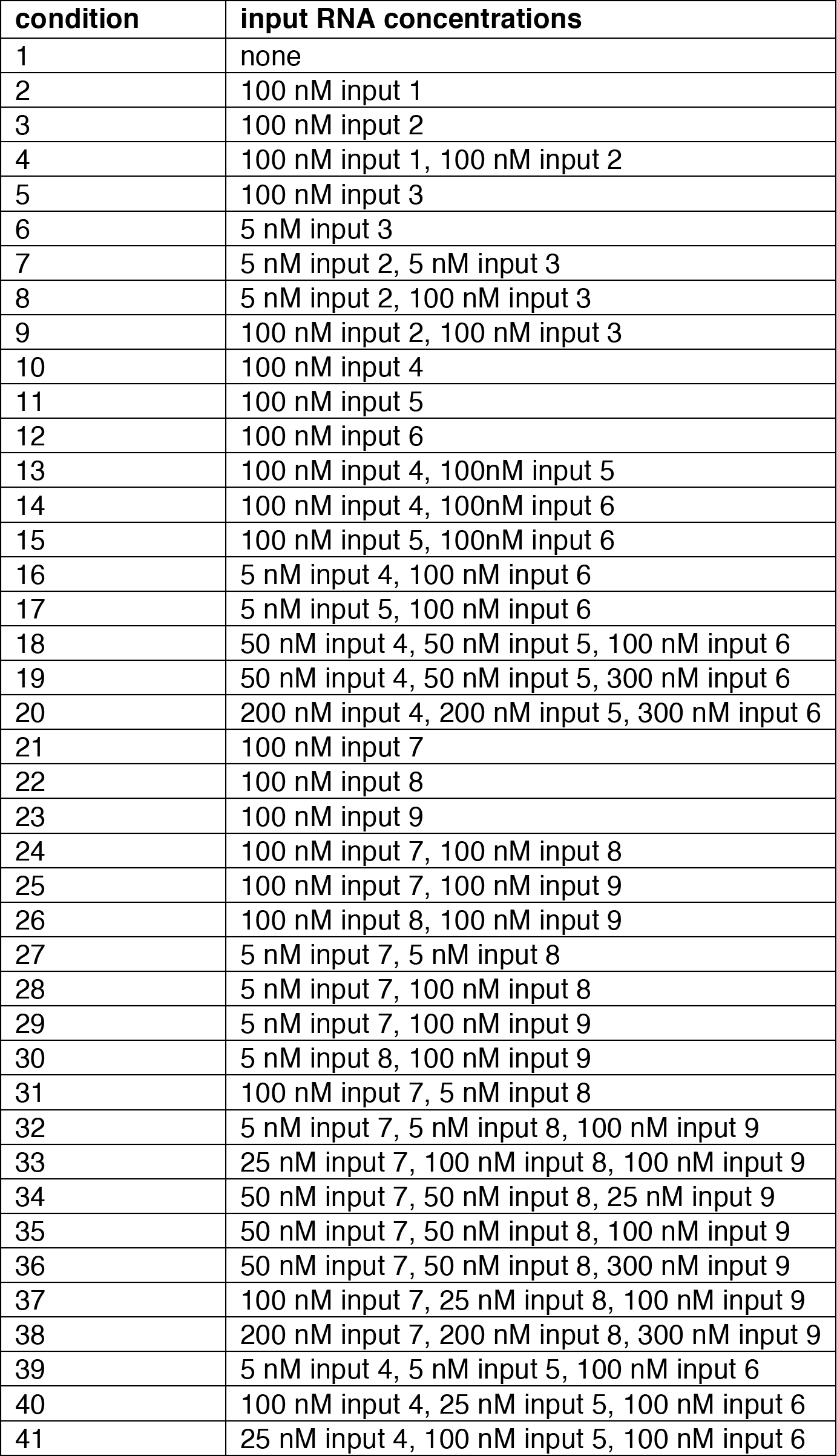
The specific concentrations of input RNAs for each experimental condition are summarized in this table.

**Table 3:**
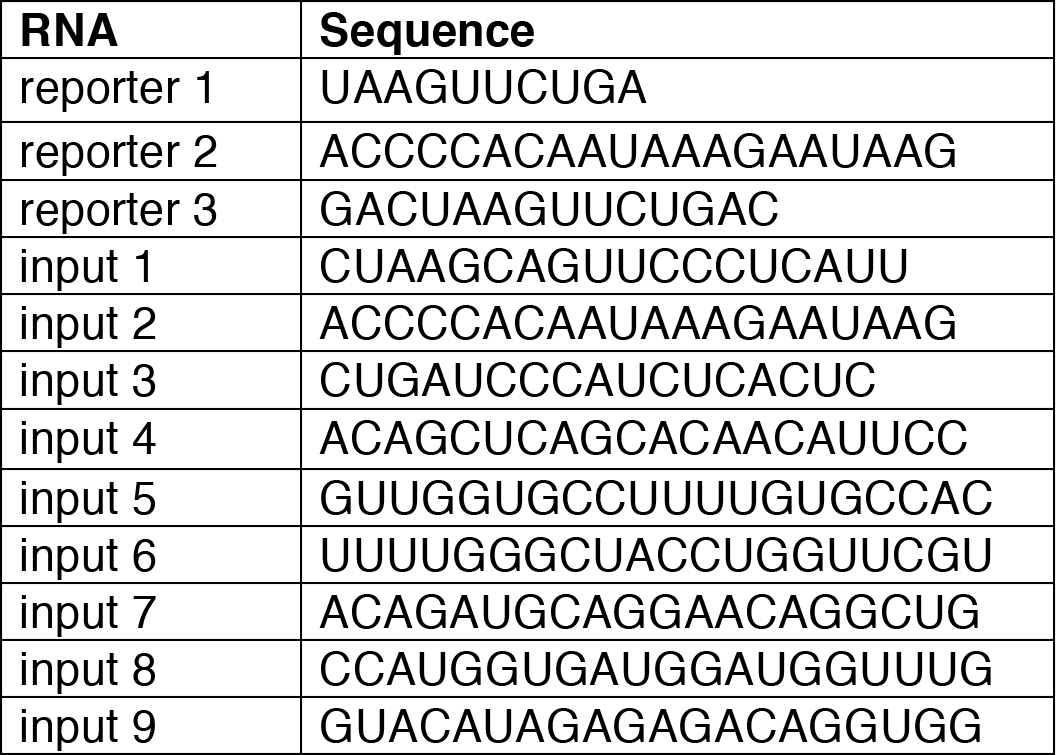
Input and reporter RNA sequences used for all puzzles are shown here.

**Table 4:**
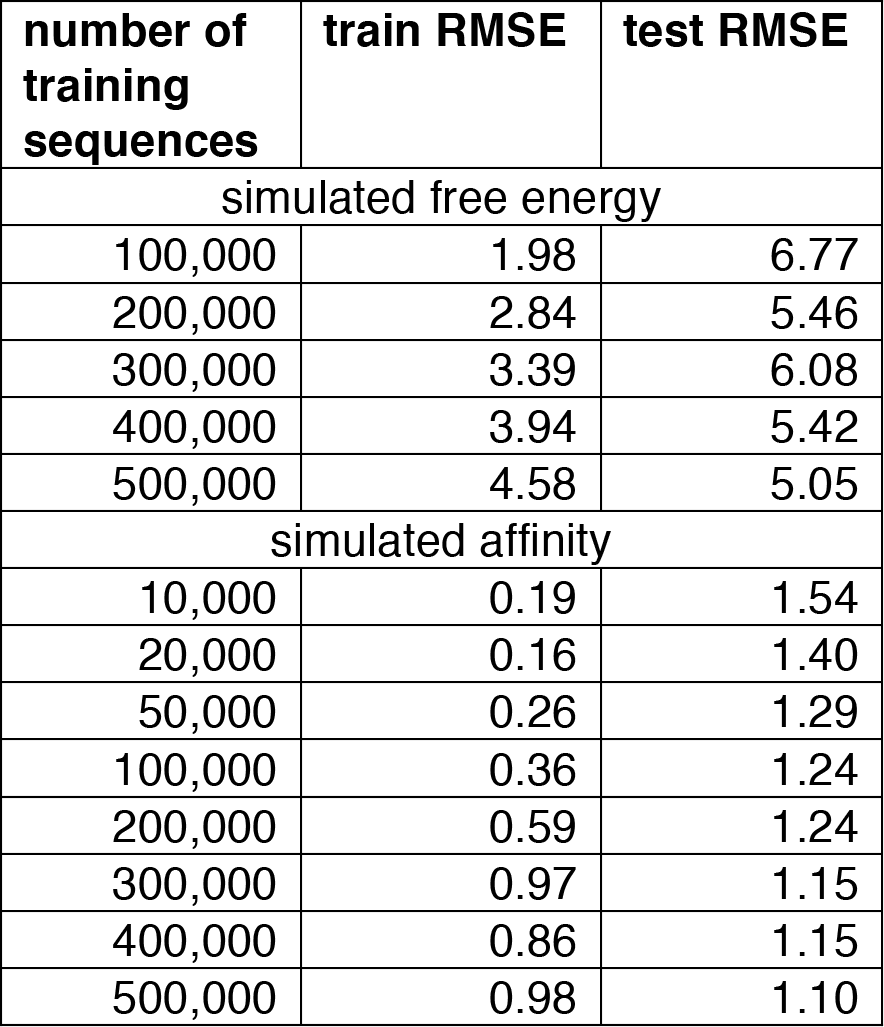
All training and test RMSEs (kcal/mol) for simulated data are summarized in this table.

**Table 5:**
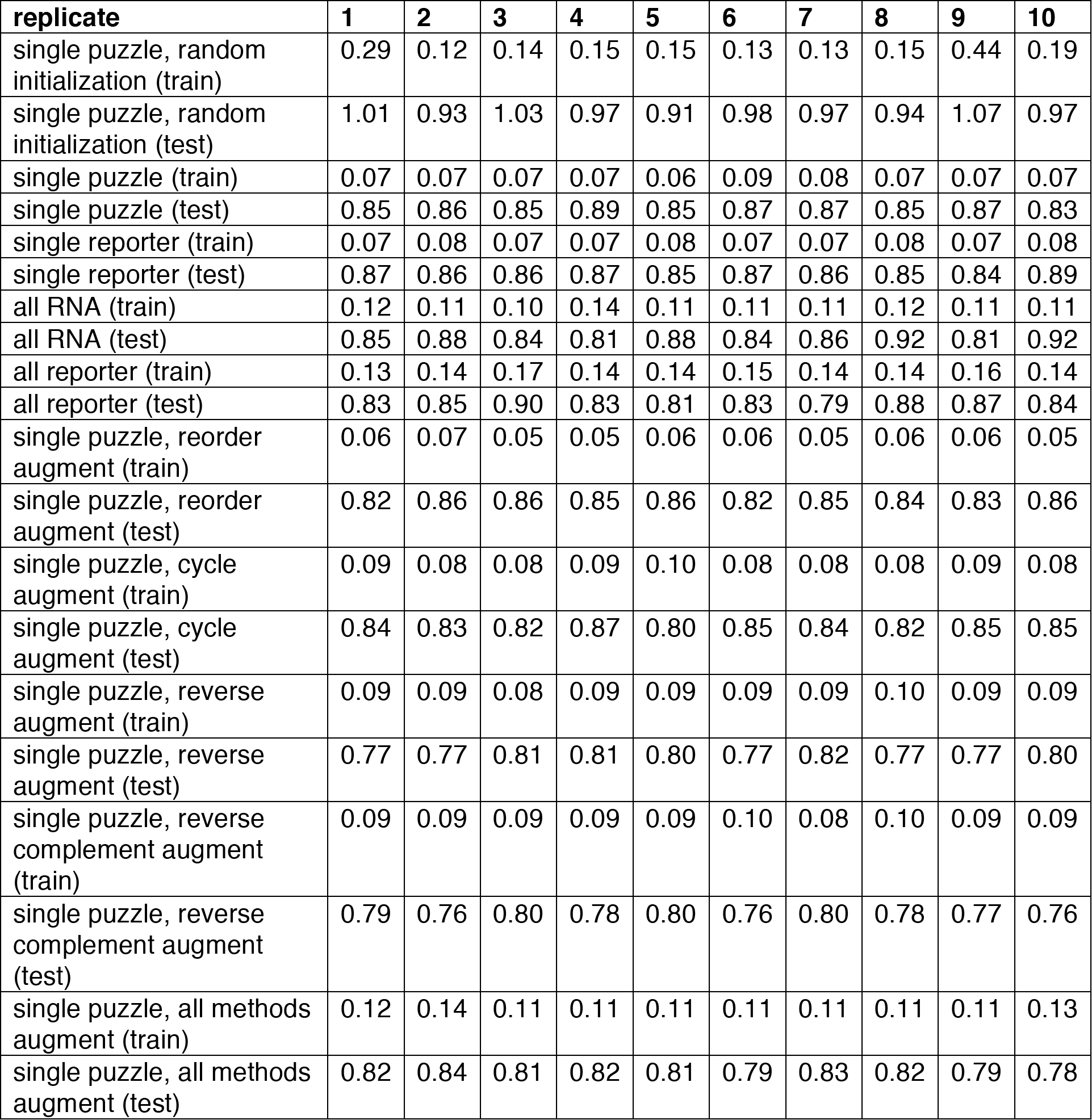
All training and test RMSEs (in kcal/mol) for experimental data are summarized in this table. All test values are filtered by a Levenshtein distance of 5 from the training data.

The median *K_d_* and *F*_max_ for each sequence variant was subsequently used and only measurements with at least 5 clusters were included in downstream analysis. For subsequent analysis, all *K_d_* measurements were converted to Δ*G* at 1M standard state for the reporter, using the expression 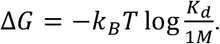

### Simulated data

Sequences were produced by generating random RNA sequences, introducing the reverse complements to the input and reporter RNA strands at random positions, and generating *n*/5 additional random mutations, where *n* is the length of the sequence. The length was chosen to be the same as those of the Eterna designs for which measurements were made. For the Turner or nearest-neighbor model, predictions were generated using the *complexes* and *concentrations* executables in NUPACK 3.0.6.[31,47] For the free energy model, the simulated free energy values were computed for the design strand along with the relevant input and reporter strands. Epistatic effects were computed by finding the difference in ΔΔ*G* between double mutants and the sum of the two single mutants:

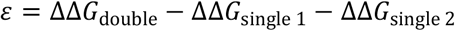

For the affinity model, complex concentrations were computed for the design strand along with the relevant input RNAs and reporter. A concentration of 1 pM was used for the design strand and 10 nM for the reporter strand, although the resulting affinity is independent of this choice as long as the reporter is in excess. These values were used to calculate the simulated Δ*G* value:

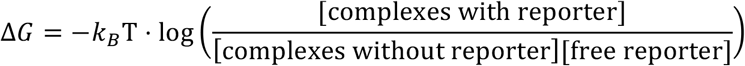

Data were generated for all 15 experimental conditions used in experimental round 9, which was used as a test set for analyses of experimental data (Table 1 & 2).

### 1-nearest neighbor model

The 1-nearest neighbor model, not to be confused with the classic Turner nearest neighbor model for RNA energetics, is a *k*-nearest neighbors model with *k*=1.[50] This model predicts the energy for a new RNA sequence as the value observed for the closest sequence in the training set. The distance metric used was Levenshtein distance between the design sequences. For input RNA concentrations not seen in the training data, the closest condition was selected based on Euclidean distance.

### Model architecture

Each measurement was identified by a design sequence, the concentrations and sequences of zero or more input RNAs, and a reporter RNA sequence. Each base in each sequence was represented as a four-element, one-hot vector. Input and design RNA vectors were multiplied by the log of their concentrations (in nM) to encode variable concentrations across conditions. For design RNA vectors, 10 µM was used as an estimate of their local concentration on the array. The input and design sequences, with each strand separated by a zero vector, served as the input to the RNN. For “single puzzle” and “single reporter” models, the sequence of the reporter was omitted from the input, as it was assumed to be the same across all data. For “all RNA” and “all reporter” models, the reporter was included as the last strand passed to the RNN (Figure S14). For the “all reporter” models that included measurements with the MS2 protein as a reporter, a fifth element was introduced into each one-hot vector to enable representation of the MS2 reporter (Figure S14). These inputs were passed through a 2-layer, 1024-unit LSTM RNN. The final state of this RNN was then processed by two fully connected layers to result in a single numerical prediction for the Δ*G* of the interaction between design and reporter strands.

### Model training

Hyperparameters were tuned using data from round 7, split into a training and validation set (Table 1). By clustering the sequences based on Levenshtein distance and splitting the resulting clusters, we ensured that similar sequences ended up in the same split of the dataset. Random search was conducted over learning rates and dropout rates to find the optimal parameter values. Various optimizers and batch sizes were also tested. All results shown were trained using the Adam optimizer, with a learning rate of 10^−^3, a dropout rate of 0.5, and a batch size of 128. All models were trained until training loss converged. With the exception of those for learning curve analysis, each model was trained 10 times with different random seeds.

For randomly initialized models, weights were initialized from a truncated normal distribution. For pretrained models, weights were initialized using the final model for the 500,000-sequence simulated dataset.

“Single puzzle” models were trained on round 7. “Single reporter” models were trained on rounds 2 and 7. “All RNA” models were trained on rounds 2–3 and 7–8. “All reporter models” were trained on rounds 1–4, 7, and 8, with rounds 5 and 6 held out as additional MS2 reporter test sets. Reported test performance was computed for round 9 unless otherwise stated. See Table 1 for details of the data collected in each round.

### Learning curves

Due to strong similarity between sequences in our dataset, clustering was used to group subsets of data instead of generating random subsets. Single-linkage hierarchical clustering was performed on the sequences to group similar designs together. The resulting tree was cut to form 10 clusters of variable numbers of sequences. The first model was then trained on only sequences in cluster 1, the second on clusters 1–2, and so on. All models were trained for 50 epochs. The test set remained the same for all models.

### Base pairing and concentration extrapolation tests

Both of these tests of whether models are biophysically realistic representations used a randomly generated design with the following sequence: CAAUCGCUCAGAACGUAGU GUACGCAGGCAGGAAUGCGGCAGAAAAGGCACUAACCGAGCCGAACCAGGUAGCCCAAA ACCUCUCC. This sequence was not used in any of the training datasets. For the base pairing test, predictions were made for all double mutants of this sequence. For the concentration extrapolation test, predictions were made for all concentration combinations of two inputs in the range from 1 pM to 1 mM.

### Design algorithm

For each round of sequence design, a random initial sequence was generated as a starting sequence. The reverse complements to the input and reporter RNA were introduced into this random starting sequence as binding sites. Each move was randomly chosen to be either a random mutation or a change in the binding sites, with equal probability. A modification to the binding sites was randomly chosen to be either a shift or change in length of the reverse complement sequence. The score was computed to be the difference between predicted affinities with and without the input RNA. The Metropolis criterion, with temperature 0.1, was applied to determine if a move was accepted or rejected. 20,000 iterations were applied to achieve the final designs. To evaluate the resulting designs, affinities were predicted using NUPACK as described above.

### Data augmentation

For the reordering augmentation dataset, *n*! data points were included for each data point in the original dataset, where *n* is the number of inputs in that experimental condition. These represent each possible ordering of the *n* input strands. For cyclic augmentation, one circular permutation of each data point was added, with the first and second halves of the design sequence swapped. For reverse augmentation, an additional copy of each data point was introduced into the augmented dataset with all sequences — inputs and design — read from 3′ to 5′ end instead of 5′ to 3′. The order of the sequences is preserved, with inputs first and design subsequently. For reverse complement augmentation, all sequences were changed to their reverse complement in an additional copy of each data point. For the combined approach, all augmentation approaches were applied to each sequence.

## Data Availability & Software

Models were implemented in Python with TensorFlow 1.0.0.[51] Data and code are available at https://github.com/wuami/ribonet.

## Acknowledgments

We acknowledge funds from the Stanford Dean’s Innovation Award (R.D.), Joint Initiative for Metrology in Biology (R.D., W.G.), NIH R01 GM100953 (R.D.), NIH R31 GM122579 (R.D.), NIH R01 GM111990 (W.G), and NSF GRFP DGE-114747 (M.W.). This work used the Extreme Science Anderson Engineering Discovery Environment (XSEDE) [52], which is supported by National Science Foundation grant number ACI-1548562.

